# Unique molecular signatures sustained in circulating monocytes and regulatory T cells in Convalescent COVID-19 patients

**DOI:** 10.1101/2022.03.26.485922

**Authors:** Andrew D. Hoffmann, Sam E. Weinberg, Suchitra Swaminathan, Shuvam Chaudhuri, Hannah Faisal Mubarak, Matthew J. Schipma, Chengsheng Mao, Xinkun Wang, Lamiaa El-Shennawy, Nurmaa K. Dashzeveg, Juncheng Wei, Paul J. Mehl, Laura J. Shihadah, Ching Man Wai, Carolina Ostiguin, Yuzhi Jia, Paolo D’Amico, Neale R. Wang, Yuan Luo, Alexis R. Demonbreun, Michael G. Ison, Huiping Liu, Deyu Fang

**Author notes:** Co-corresponding authors. Huiping Liu, MD, PhD, Northwestern University, 303 E Superior St, Chicago, IL 60611. Deyu Fang, MD, PhD, Northwestern University, 300 E Chicago St, Chicago, IL 60611. Michael Ison, MD, Northwestern University, 645 North Michigan Avenue Suite 900, Chicago, Illinois 60611. Co-first authors with equal contributions (Andrew D. Hoffmann, Sam E. Weinberg).

## Abstract

Over two years into the COVID-19 pandemic, the human immune response to SARS-CoV-2 during the active disease phase has been extensively studied. However, the long-term impact after recovery, which is critical to advance our understanding SARS-CoV-2 and COVID-19-associated long-term complications, remains largely unknown. Herein, we characterized multi-omic single-cell profiles of circulating immune cells in the peripheral blood of 100 patients, including covenlesent COVID-19 and sero-negative controls. The reduced frequencies of both short-lived monocytes and long-lived regulatory T (Treg) cells are significantly associated with the patients recovered from severe COVID-19. Consistently, sc-RNA seq analysis reveals seven heterogeneous clusters of monocytes (M0-M6) and ten Treg clusters (T0-T9) featuring distinct molecular signatures and associated with COVID-19 severity. Asymptomatic patients contain the most abundant clusters of monocyte and Treg expressing high CD74 or IFN-responsive genes. In contrast, the patients recovered from a severe disease have shown two dominant inflammatory monocyte clusters with S100 family genes: S100A8 & A9 with high HLA-I whereas S100A4 & A6 with high HLA-II genes, a specific non-classical monocyte cluster with distinct IFITM family genes, and a unique TGF-β high Treg Cluster. The outpatients and seronegative controls share most of the monocyte and Treg clusters patterns with high expression of HLA genes. Surprisingly, while presumably short-ived monocytes appear to have sustained alterations over 4 months, the decreased frequencies of long-lived Tregs (high HLA-DRA and S100A6) in the outpatients restore over the tested convalescent time (>= 4 months). Collectively, our study identifies sustained and dynamically altered monocytes and Treg clusters with distinct molecular signatures after recovery, associated with COVID-19 severity.

## Introduction

The COVID-19 pandemic caused by SARS-CoV-2 has infected more than 298 million people and caused over 5.4 million deaths since its emergence in late 2019(1-3). New, evolving variants of concern and delays in vaccination (over 90% unvaccinated in low-income countries) mean that it will continue to disrupt the global society for extended time in the future. Shortly after COVID-19 became a global pandemic, post-acute health issues were reported(4). Typical issues included shortness of breath, muscle fatigue, and prolonged loss of smell. Other reported symptoms include lingering headaches and cognitive difficulties, skin rashes and gastrointestinal discomfort. Combined, these constellation symptoms are now informally referred to as “long COVID” and likely represent a diverse group of post-infectious syndromes(5-7). These symptoms have been reported to occur at higher frequency in patients with severe initial infection(8, 9). In addition to these long-term ailments, COVID has frequently been characterized by high amounts of inflammation. Together, these suggest that long-term changes in the immune system even following recovery from COVID could be at the root of some of these long-term symptoms.

An efficient immune response against invading pathogens including SARS-CoV-2 requires the early activation of innate immunity, a nonspecific but quick frontline response able to control infection(10-14). This effective innate immune response plays a critical role to mount the antigen-specific adaptive immunity. The latter contributes to clearing the infection and preventing reinfection by the same pathogen and more importantly, often sustains the memory response ready for future threatens by the same pathogen(15). Numerous examinations of patient response to COVID-19 at the single-cell level have been performed since the disease became widespread. Multiple meta-analyses of single-cell RNA-sequencing data sets have defined the several immune dysregulations are involved in COVID-19 pathogenicity including lymphopenia, impaired IFN response, hyperactivation of myeloid cells and dysregulated macrophage and monocyte functions(11, 16-26). A common feature of SARS-CoV-2 infection in severe COVID-19 patients is lymphopenia with a drastic reduction in both T and B lymphocytes in the circulating blood. This lymphopenic response is often negatively associated with the viral load of SARS-CoV-2 as well as the disease severity(21, 27). In addition, Wilk et al(21) performed some of the earliest single-cell RNA sequencing of COVID-19 patient-derived cells, identifying changes in the myeloid compartment as a consequence of more severe disease states. Despite these progresses, critical questions regarding to whether the dysregulated immune cell phenotypes observed during SARS-CoV-2 infection persist in COVID-19 patients after their full recovery, and if yes, how long the fingerprints in gene expression can be maintained, remain to be answered. In order to understand the post-COVID immune response, we utilized the single cell RNA sequencing approach and determined the molecular signatures of monocytes and understudied regulatory T (Treg) cells in the circulating blood from convalescent COVID-19 patients (when they were unvaccinated). Our studies revealed several unique clusters and molecular signatures in both populations sustained during the recovery phase after SARS-CoV-2 infection.

## Results

### Association of monocyte and Treg populations with COVID-19 pathogenicity

During the convalescent phase following SARS-CoV-2 infection, the immune system consists of two distinct cellular pools, the virus experienced and naïve cells. As a patient recovers, the SARS-CoV-2 experienced pool changes from a mixture of short lived myeloid, lymphoid and tissue resident cells to predominantly long-lived lymphocytes and tissue resident macrophages. Thus, in convalescent patients, the circulating monocytes and granulocytes are predominantly COVID-19 naïve. To assess whether and how SARS-CoV-2 infection maintains a sustained impact to immune function, we characterized the levels of circulating monocytes from convalescent COVID-19 patients. We enrolled 100 convalescent patients at least three weeks after recovery from COVID-19 and analyzed both plasma IgG antibodies specific to SARS-CoV-2 receptor binding domain (RBD) and cellular immunity profiling (**Table 1, Figure 1A**). Patients were classified into 5 groups based on severity of disease and RBD-IgG levels (**Supplementary Figure 1B**)(28, 29), including sero-negative healthy control group (undetectable RBD-IgG and PCR negative/no PCR), asymptomatic (detectable RBD-IgG but symptom free during SARS-CoV-2 infection), outpatient (mild symptoms, detectable RBD-IgG), and hospitalized groups (non-ICU and ICU subgroups). As expected, severe COVID symptoms in hospitalized groups (non-ICU and ICU) correlated with higher IgG levels (**Supplementary Figure 1A**). Consistantly, plasma capacity to block RBD binding to ACE2-expressing cells was positively correlated with RBD-IgG levels and disease severity (**Supplementary Figure 1B**). While we expected that time from disease onset would likewise correlate due to eventual senescence of antibody producing cells, the plasma RBD-IgG levels and capacity to inhibit RBD binding in outpatients gradually decline by four months after their recovery (**Supplementary Figure 1C-D**).

**Table 1.**

|  | Sero-negative<br>(N=5) |  | Asymptomatic<br>(N=18) |  | Outpatients<br>(N=56) |  | Hospitalized<br>(N=12) |  | ICU<br>(N=9) |  |
| --- | --- | --- | --- | --- | --- | --- | --- | --- | --- | --- |
|  | mean /<br>count | SD<br>/ % | mean /<br>count | SD<br>/ % | mean /<br>count | SD<br>/ % | mean /<br>count | SD<br>/ % | mean /<br>count | SD<br>/ % |
| Age, years | 39.4 | 15.7 | 33.4 | 8.2 | 42.3 | 15.4 | 48.6 | 11.5 | 50.7 | 16.5 |
| Sex, male | 0 | 0.0% | 9 | 50.0% | 22 | 39.3% | 5 | 41.2% | 6 | 66.7% |
| Race |  |  |  |  |  |  |  |  |  |  |
| Black | 0 | 0.0% | 0 | 0.0% | 4 | 7.1% | 6 | 50.0% | 1 | 11.1% |
| White | 3 | 60.0% | 8 | 44.4% | 30 | 53.6% | 5 | 41.7% | 8 | 88.9% |
| Asian | 0 | 0.0% | 4 | 22.2% | 1 | 1.8% | 0 | 0.0% | 0 | 0.0% |
| Other | 2 | 40.0% | 6 | 33.3% | 21 | 37.5% | 1 | 8.3% | 0 | 0.0% |
| SOFA<br>score* | NA | NA | NA | NA | NA | NA | NA | NA | 6.4 | 3.5 |
| Intermediate to advanced interventions |  |  |  |  |  |  |  |  |  |  |
| Vasopressor use | 0 | 0.0% | 0 | 0.0% | 0 | 0.0% | 1 | 8.3% | 7 | 77.8% |
| High flow nasal cannula | 0 | 0.0% | 0 | 0.0% | 0 | 0.0% | 0 | 0.0% | 5 | 55.6% |
| Non-invasive ventilation | 0 | 0.0% | 0 | 0.0% | 0 | 0.0% | 0 | 0.0% | 1 | 11.1% |
| Mechanical ventilation | 0 | 0.0% | 0 | 0.0% | 0 | 0.0% | 0 | 0.0% | 7 | 77.8% |
| LOS, days | NA | NA | NA | NA | NA | NA | 6.8 | 5.3 | 29.6 | 19.2 |
| Onset to sampling, days | NA | NA | 80.7 | 51.4 | 82.6 | 51.1 | 145.1 | 53.8 | 119.6 | 39.1 |

**Figure 1:**
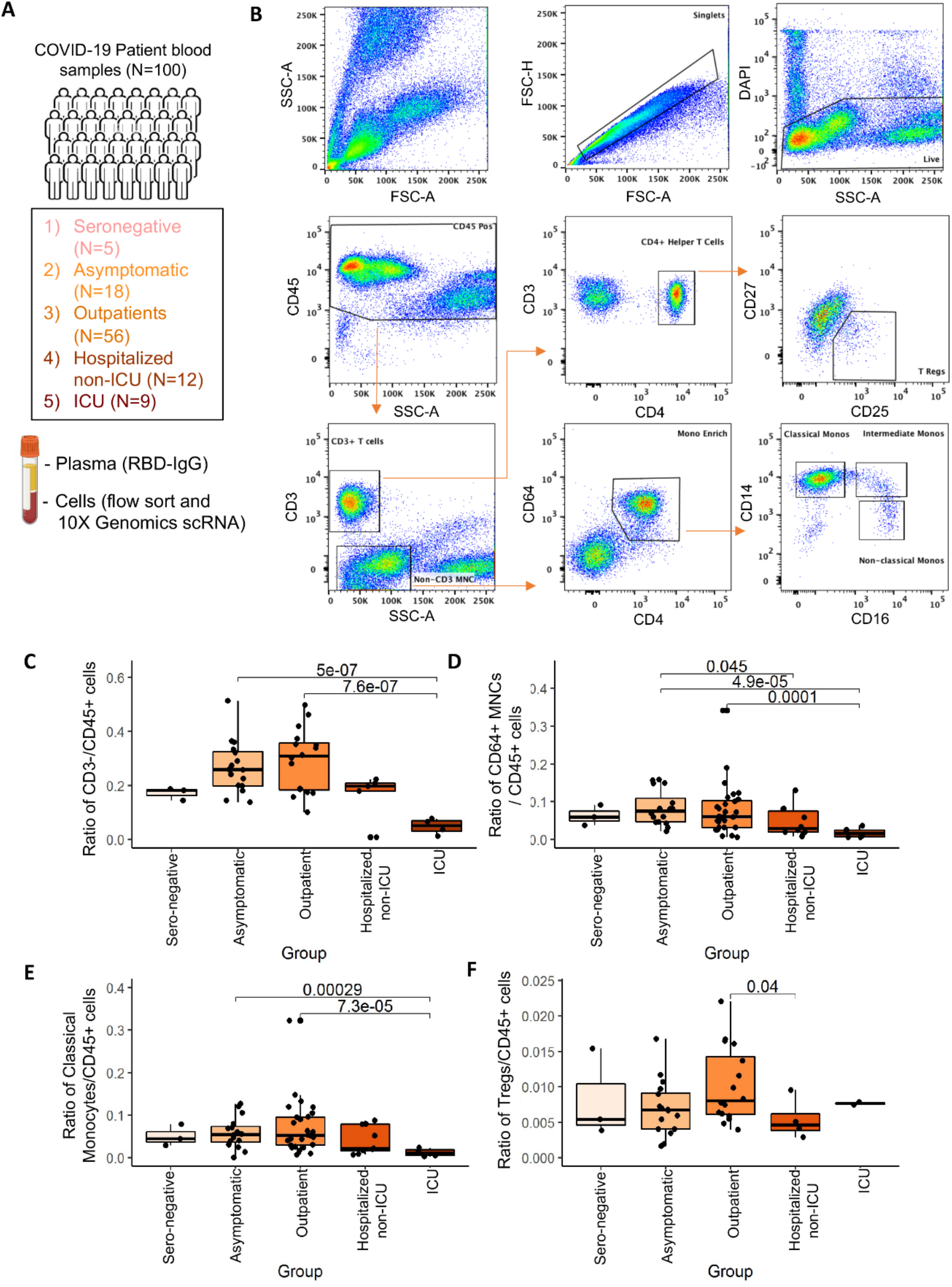
Flow profiles of circulating immune cells associated with COVID-19 patients(N=100). **(A)** Number of patients enrolled and cell isolation strategies for RBD-IgG, flow and single-cell sequencing analysis. **(B)** Representative flow cytometry analysis showing the gating and sorting strategies for WBCs isolated from convalescent COVID-19 patients. CD45^+^DAPI^-^ alive cells were gated into CD3^+^ and CD3^-^ within the SSC^low^ groups. CD3^+^ cells were further sub-divided into CD4^+^ Helper T cells, and Treg cells were sorted as CD4^+^CD25^high^CD127^low^ population. CD3-cell fraction were sorted based on CD4^dim^CD64+ MNCs fraction (enriched for monocytes); these cells were further sub-divided into CD14^+^CD16^-^ classical monocytes, CD14^-^CD16^+^ nonclassical, and CD14^+^CD16^+^ intermediate monocytes for analysis. **(C-E**) Ratio of flow gated CD3^-^ mononuclear (B), CD64^+^ monocytes (C), and CD14^+^ classical monocytes (D) within CD45^+^ WBCs among five groups of COVID-19 patients (sero-neg, asymptomatic, outpatients, hospitalized-no ICU, ICU), with elevation in mild disease groups (asymptomatic/outpatient) in comparison with severe ICU patients. (**F**) Ratio of flow gated Treg cells within CD45^+^ WBCs among five groups of COVID-19 patients, elevated in outpatients in comparison with hospitalized patients. Boxplots in B-E indicate quartiles. Comparison p-values calculated using Wilcoxon signed rank test.

Initial flow cytometric analysis of the gated CD45^+^ peripheral white blood cells (WBCs) identified monocytes based on the smaller cell size and lower granularity than granulocytes, negative for expression of CD3, dim positive for CD4, and positive for CD64, a specific marker for human myeloid cells, particularly macrophages and monocytes (**Figure 1B**). These CD45^+^CD3^-^ CD4^dim^CD64^+^ cells were further sub-divided based on their expression of CD14 and CD16 into: CD14^+^CD16^-^ classical monocytes, CD14^+^CD16^+^ intermediate monocytes, and CD14^-^CD16^+^ non-classical monocytes (**Figure 1B**). Interestingly, the frequency of monocyte population showed a dynamic pattern in association with the COVID-19 disease severity. As shown in **Figure 1C-E**, we observed a statistically significant increase in all monocytes and the classical CD14^+^ monocyte levels relative to total CD45^+^ cells in recovered COVID-19 patients with less severe disease (asymptomatic and outpatient), which are found at lower levels in patients recovered from the most severe disease (ICU). Therefore, the decline in monocyte populations during the recovery phase appear to be a signature memory for severe disease.

One possible explanation for the long-term changes seen in circulating monocytes in patients recovered from COVID-19 is global alterations to the immune system driven by long-lived adaptive immune cells. CD4^+^CD25^+^FoxP3^+^ Tregs are known to modulate broad aspects of the immune response including maintenance of immune tolerance and global immunosuppression, and we hypothesized that changes in the Treg compartment may persist following the recovery from COVID-19. To assess changes in Treg population in convalescent patients, CD45^+^CD3^+^ T lymphocytes were further analyzed by CD4 expression as helper T cell population. Tregs were then defined as CD4^+^CD25^+^CD127^low^ populations as reported (30) (**Figure 1B**). Importantly, the Treg cell frequency during the convalescent phase was increased in less severe outpatients compared to hospitalized patients (**Figure 1F**), suggesting that SARS-CoV-2 infection may promote an immunosuppressive state in patients with mild disease.

### Identification of persistent subsets of monocytes with distinct molecular signatures from convalescent COVID-19 patients

To further investigate the effect of COVID-19 severity on the molecular signature of convalescent monocyte phenotypes, we sorted CD45^+^CD3^-^CD64^hi^CD4^dim^ intermediate side scatter cells from 50 convalescent patients and sero-negative controls, and subjected to the 10X Genomics single cell sequencing (See **Supplementary Table S1** for further demographic information). Clustering analysis identified seven distinct clusters, each of which is specified with two top enriched genes, including five CD14^+^ classical monocyte clusters and one distinct CD16^+^ non-classical monocyte cluster M5_ISG15_IFI6, and one dendritic cell cluster M6_ZEB2_CCNL1 showing lower expression of both HLA and S100 family genes (**Figure 2A-B**). Further evaluation of the five classical monocyte subsets found five subsets with inflammatory features. Clusters M1_S100A8_S100A9 and cluster M3_S100A6_S100A4 both showed high but distinct S100 genes (cluster 1 with high levels of S100A8 & 9, but cluster 3 with high S100A6 & 4). Also, the M1_S100A8_S100A9 monocytes show uniquely high levels of class I HLA molecules compared to cluster M3_S100A6_S100A4, which expressed predominantly HLA class II genes (**Figure 2C-D**). Cluster M0_HLA-B_EEF2 and cluster M2_CD74_HLA-DR shared relatively similar gene expression patterns but Cluster M0 displayed enhanced S100A8/9, STAT1, and ZEB2 genes within multiple high level heatmap blocks genes and appear to be intermediate populations (**Figure 2C & D**). Consistant with previous studies showing the altered IFN response during the acute SARS-CoV-2 infection is responsible for severe disease (13, 31), we detected two clusters with high but distict IFN responsive genes: the classical monocyte cluster M5_ISG15_IFI6 displayed a unique gene expression pattern characterized by high expression of interferon stimulated genes including ISG15, IFI6, IFIT3, and IFIT2. In contrast, the cluster M4_IFITM_FCGR3A, which belong to CD16^+^CD14^-^ non-classical monocytes, expressed high levels of different IFN responsive genes in particular the Interferon Induced Transmembrane Protein (IFITM) family genes (IFITM1-3) (**Figure 2C & D**).

**Figure 2.**
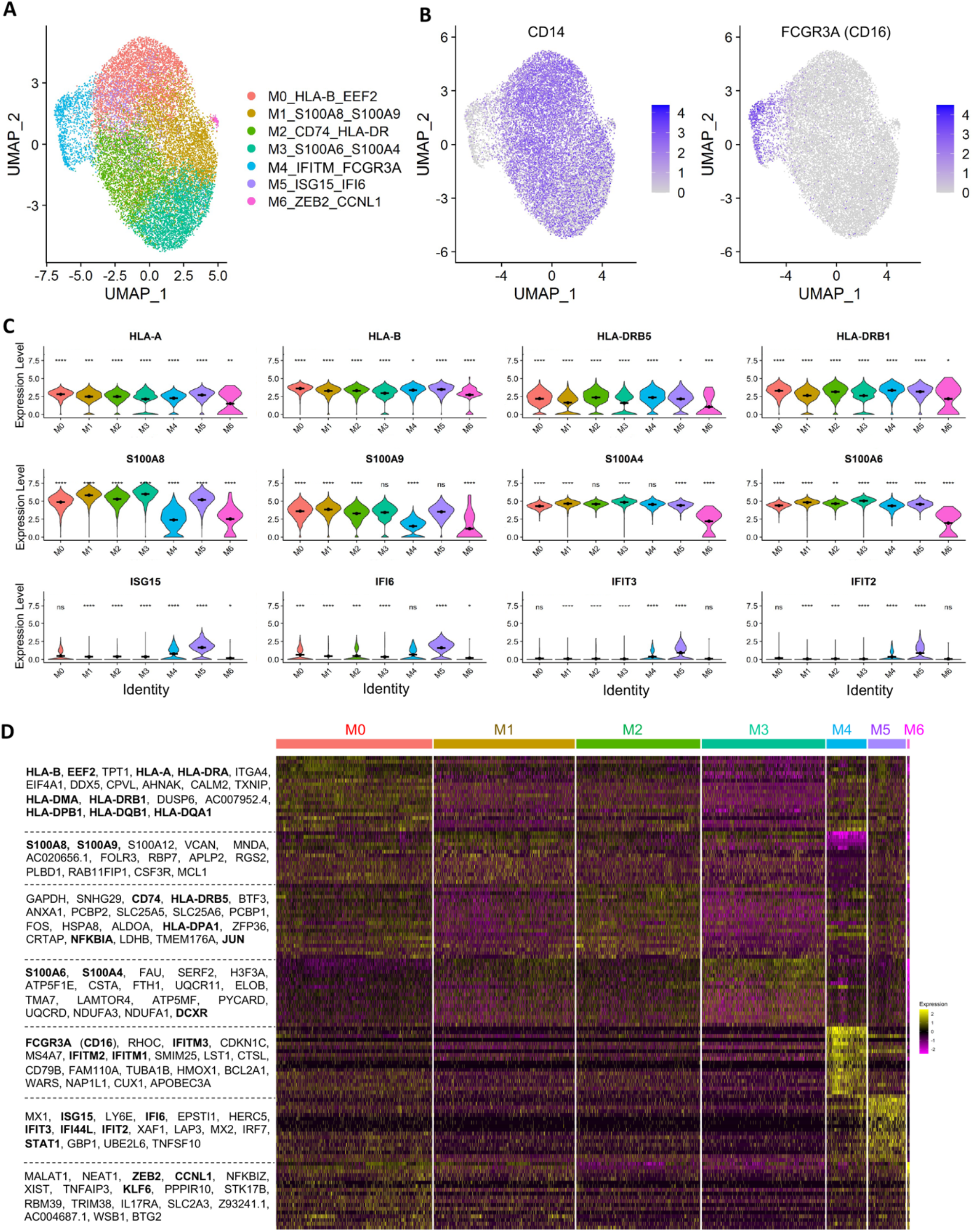
Analysis of the sustained molecular signatures in monocytes from convalescent COVID-19 patients. **(A)** Dimensional UMAP plot of monocytes from convalescent COVID-19 patients and sero-negative controls. **(B)** Expression of CD14, indicating classical monocytes, and FCGR3A/CD16, indicating nonclassical monocytes, on the UMAP projection from A. Classical monocytes make up the majority of the cells and five of the seven clusters. Nonclassical monocytes are principally cluster M4 on the dimensional plot in A. **(C)** Expression of selected genes across clusters. Class I HLA molecules (HLA-A and HLA-B) are strongly expressed in cluster M0, while class II HLA molecules (HLA-DRB5 and HLA-DRB1) are more strongly expressed in cluster M2. S100 genes (second row) are most highly expressed in clusters M1 and M3. Interferon-responsive genes (third row) are most present in cluster M5, which appears to be a unique interferon-responsive cluster of monocytes. **(D)** Heatmap showing the expression levels of selected genes across all cells. Nonclassical monocyte associated genes are clearly visible in cluster M4, as are interferon-responsive genes in cluster M5. Significance of gene expression difference in individual clusters evaluated by Wilcoxon comparison of cluster-cell expression levels with all-cell expression levels. Significance values: ns: p>0.05, * p<0.05, ** p<0.01, *** p<0.001, **** p<0.0001

Importantly, these monocyte clusters with distinct molecular signatures were found to be associated with COVID-19 severity. The S100 family gene-high cluster M1-S100A8_S100A9 decreased but the immune regulatory receptor CD74-high cluster M2-CD74_HLA-DR increased in the convalescent patients with asymptomatic COVID-19 vs other patient groups. In addition, the cluster M5_ISG15_IFI6 with high in IFN signature genes were enriched in asymptomatic, but not other COVID-19 patients with severe disease (**Figure 3A**). Of note, in outpatients the M5_ISG15_IFI6 cluster with IFN response genes was lost during the early (0-1 month) but restored later (2 months or after) disease onset (**Figure 3B, Supplementary Table S2**). IFN response gene IFI6 was highly enriched in the monocytes isolated from asymptomatic patients and the patients during 2-3 months after infection, whereas ISG15 was mostly depleted in the monocytes of ICU patients **(Figure 3C-D, Supplementary Table S2**). It has been well established that the impairments in the IFN response is the key critical for anti-SARS-CoV-2 immune response(13, 31), our data imply a possibility that the patients with a better expansion of monocytes in M5_ISG15_IFI6, as well as Clusters M2_CD74_HLA-DR, have a favorable clinical outcome. Consistent with this notion, further analysis reavealed a trend in IFN-responsive monocyte reduction during the early recovery phase of outpatients, which gradually returned 4 months after acute infection (**Figure 3D**).

**Figure 3.**
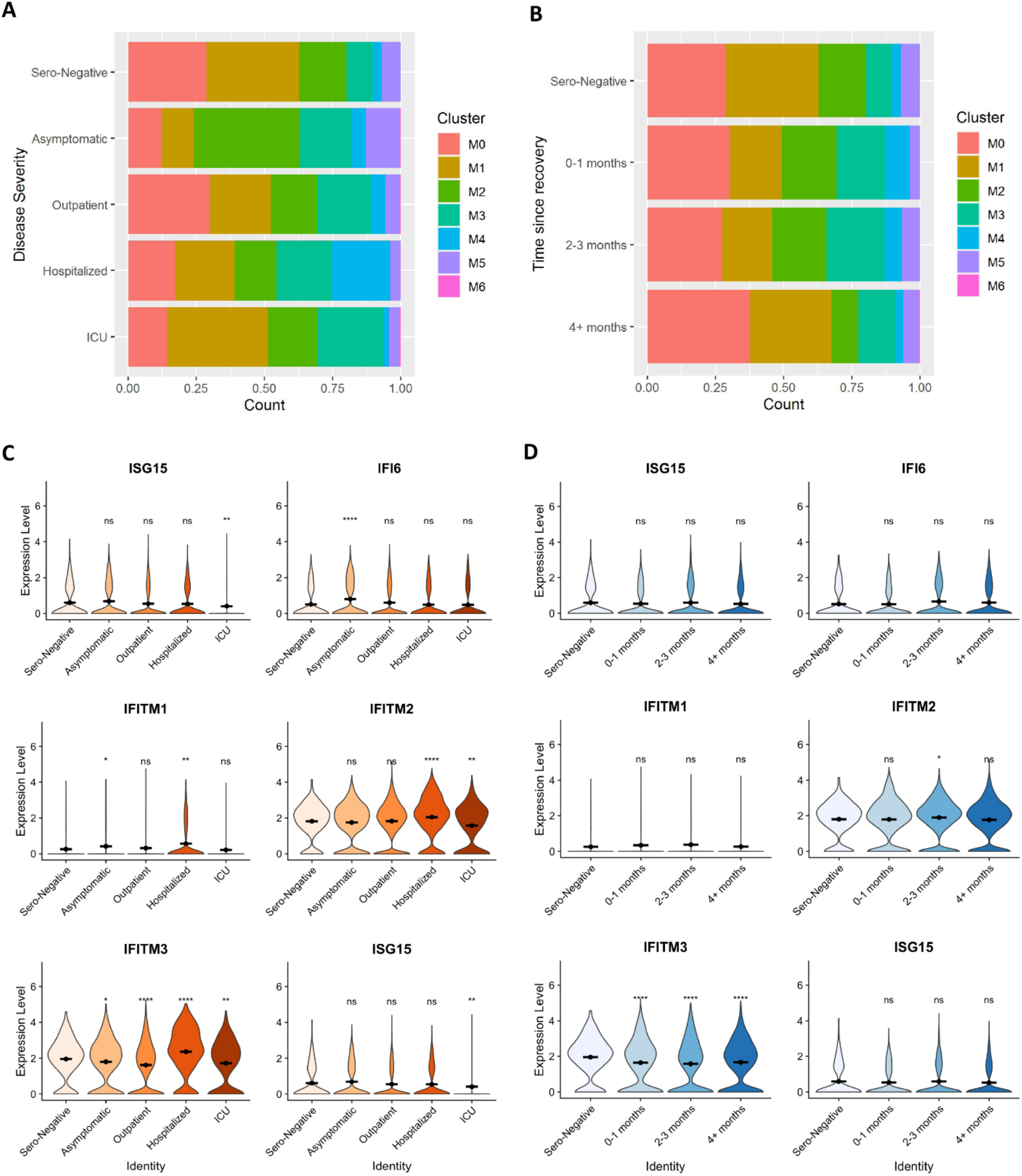
Association of monocytes’ signatures with COVID-19 disease severity and recovery. **(A)** Proportion of monocyte cells from each disease severity category in each cluster. Cluster M5 (interferon-stimulated monocytes) appears enriched in asymptomatic patients, while cluster M4 (nonclassical monocytes) appears enriched in hospitalized patients. **(B)** Proportion of monocytes in outpatients and sero-negative controls separated by time. Changes in monocyte proportions in these patients are largely not sensitive to time since disease. **(C)** Interferon stimulated gene expression in cells separated by disease severity. Both genes appear to be most highly expressed among asymptomatic patients. **(D)** Expression of interferon stimulated genes in cells derived from outpatients and sero-negative controls only separated by time from disease. These genes show very little change over time. C-D: Significance of gene expression difference in individual groups evaluated by Wilcoxon comparison of group-cell expression levels with Sero-Negative-cell expression levels. Significance values: ns: p>0.05, * p<0.05, ** p<0.01, *** p<0.001, **** p<0.0001

Along with changes in the classical monocyte populations, the non-classical cluster M4_IFITM_FCGR3A appeared to be enriched specifically in recovered hospitalized patients (**Figure 3A**). Importantly, when examined in outpatients this enrichment was time dependent as the proportion of non-classical monocytes was enriched during the early recovery phase (0-1 month) since acute infection, which gradually returned to a comparable level as that in sero-negative controls (**Figure 3B**). Interestingly, unlike the classical monocyte cluster M5_ISG15_IFI6, the non-classical monocytes in Cluster M4_IFITM_FCGR3A/CD16 have an dynamic changes in expression of distinct interferon-induced genes IFITM1-3, which are specifically enriched in the hospitalized non-ICU group. In contast, their levels, in particular IFITM3, in outpatients were significantly reduced **(Figures 2D, 3A & 3C)**. Further longitutional analysis show the reduction in IFNTM3 levels sustained for morethan 4 months after COVID-19 recovery **(Figure 3D)**. The IFITM proteins have been identified as cell-autonomous proteins that suppress the early stages of viral replication(32), our data suggest that the increase of this non-classical cluster might be responsible for the severe COVID-19 pathogenesis. Circulating, classical monocytes are known to have a short lifespan of no more than a few days, while non-classical monocytes display longer survival, but still are unlikely to live for more than 1 or 2 weeks (33). Thus, the monocytes we observed in our analysis likely developed following clearance of SARS-CoV-2 and the restoration of some degree of immune homeostasis in these patients. In that context, our data demonstrated the long-term changes in monocyte phenotypes in patients recovered from COVID-19 is surprising, which suggests that acute SARS-CoV-2 infection induces immune system alteration even after elimination of replicating virus.

### scRNA-Seq identifies unique Treg clusters and long lasting Treg cell signatures in patients recovered from COVID-19

Our discovery that the frequency of FoxP3^+^ Tregs is increased in the circulating blood from recovered COVID-19 outpatients but decreased in hospitalized patients (**Figure 1F**), suggesting that dynamic changes in circulating Treg frequency may impact both acute COVID-19 pathogenesis and long-term complications. We sorted the CD45^+^CD3^+^CD4^+^CD25^high^CD127^low^ population, which are known to sufficiently define human Tregs from PBMC (30), from 27 convalescent COVID-19 patients and sero-negative controls (**Supplemental Table S3** for further information), and subjected them to the 10X Genomics single cell sequencing platform. Clustering analysis identified 10 clusters (**Figure 4A**), indicating that Tregs in the circulating blood from both COVID-19 patients and healthy donors are highly heterogeneous.

**Figure 4.**
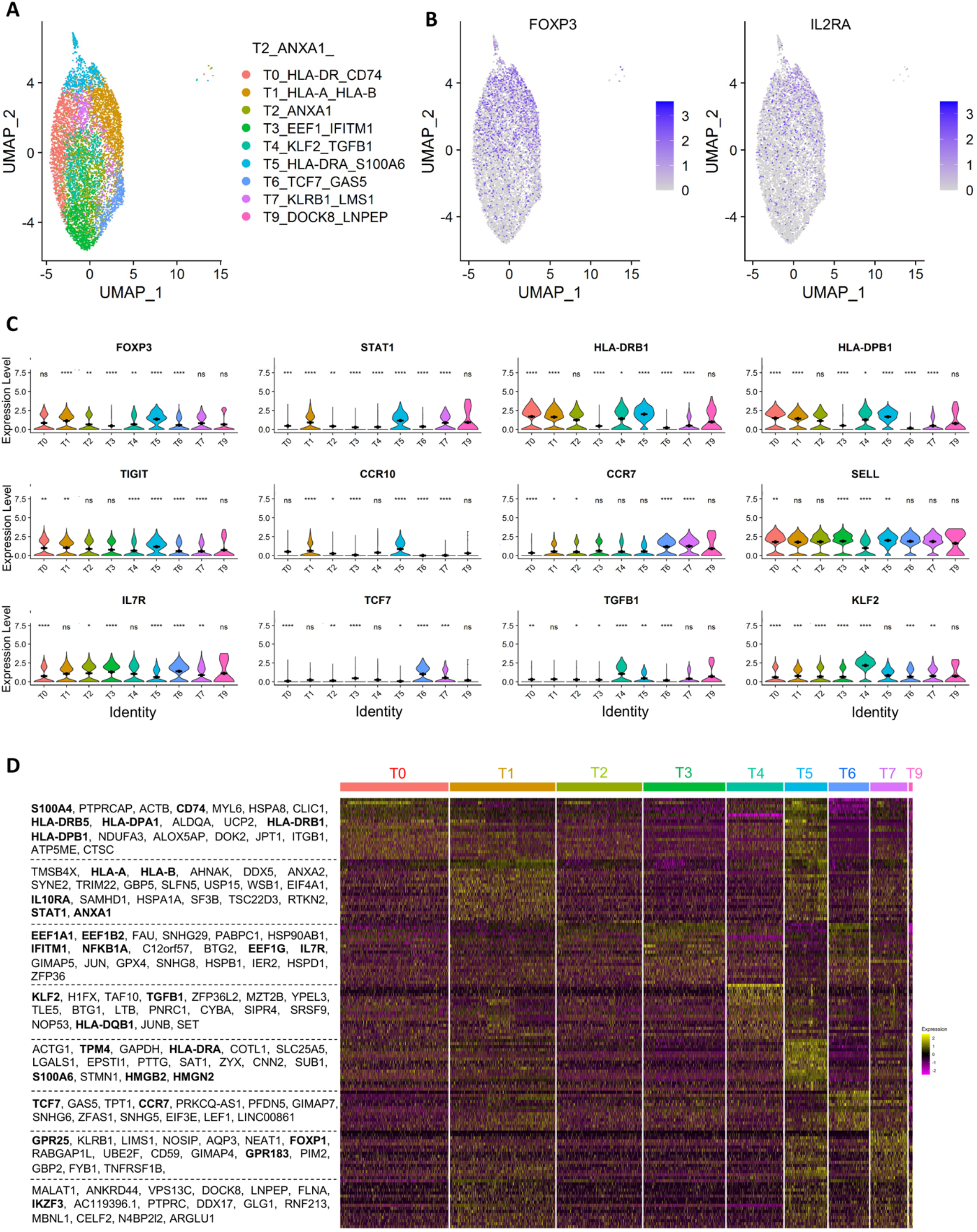
Analysis of the sustained Treg molecular signatures from convalescent COVID-19 patients. **(A)** Dimensional UMAP plot of Tregs from convalescent COVID-19 patients and sero-negative controls. **(B)** Expression of FOXP3 and CD25/IL2RA on the UMAP projection from A. **(C)** Expression of selected genes across clusters. FOXP3 and STAT1 are strongly expressed in clusters T1, T5 and T7, indicating effector cells. Clusters T2, T3, T6 and T9 show lower levels of FOXP3 and STAT1 but higher levels of SELL and IL7R, indicating central/homeostatic Treg cells. Cluster T5 expresses high levels of CCR10, CCR7 and FOXP3, potentially indicating a tissue-migratory effector Treg phenotype. Cluster T4 is unique to patients recovering from COVID-19 and expresses high levels of TGFB1 and KLF2. **(D)** Heatmap showing the expression levels of selected genes across all cells. Significance of gene expression difference in individual clusters evaluated by Wilcoxon comparison of cluster-cell expression levels with all-cell expression levels. Significance values: ns: p>0.05, * p<0.05, ** p<0.01, *** p<0.001, **** p<0.0001

Further annotation of gene expression of the individual clusters identified related signatures between clusters, and each cluster is defined by one or two top enriched genes. Treg clusters T1_HLA-A_HLA-B and T5_HLA-DRA_S100A6 displayed high expression of FOXP3 and STAT1, which are consistent with an activated/effector Treg phenotype. To support this designation, clusters T1 and T5 also express high levels of HLA class II molecules and TIGIT (**Figure 4B-D**), which is thought to define a unique Treg subset that preferentially suppresses Th1 and Th17-driven inflammatory responses(34). Tregs in T1_HLA-A_HLA-B and T5_HLA-DRA_S100A6 also show a tissue migratory Treg-signature with high expresses levels of CCR10, CCR7, and FOXP1. In contrast, Tregs in clusters T2_ANXA1 and T3_EEF1_IFITM1, have reduced expression of TIGIT and class two HLA molecules. In addition, the cluster T6_TCF7_GAS5 shows lower levels of FOXP3 and CD25, as well as HLA family genes, but displays higher levels of central/homeostatic Treg markers including TCF7, SELL and IL-7R. Cluster T7_KLRB1_LMS1 also showed increased expression of CCR7, TCF7 and KLF2 consistent with a less suppressive Treg phenotype. Interestingly, a Treg population (cluster T4_KLF2_TGFB1) with elevated expression of TGF-β and KLF2 was only identified in a subset of covelesent COVID-19 patients, which was not identified in sero-negative patients (**Figure 4B & 4C**). The clinical significance of this cluster T4_KLF2_TGFB1 population is unclear. However, as Treg-specific production of TGF-β is known to have a variety of immunologic roles, the fact that this population is only observed in patients recovering from COVID-19 suggests a possible role in COVID-19 pathogenesis and recovery. Importantly, because this cluster was only identified in patients who had recovered from SARS-CoV-2 infection, these cells may identify a distinct SARS-CoV-2-associated immunological profile. In addition to these 7 clusters with unique gene expression signatures, Tregs in cluster 0, T0_HLA-DR_CD74, share a largely common gene expression profile with all the rest clusters, implying a possibility that Tregs in this cluter are a intermediate population. Tregs in cluster 9, T9_DOCK8_LNPEP, which are rather minor, show reduced expression of genes enriched in cluster T3_EEF1_IFITM1 including EEF family and IFN responsive genes **(Figure 4D)**.

Similar to monocytes in convalescent COVID-19 patients, the relative proportions of the identified Treg clusters show differences that are correlated with both initial disease severity and the time post recovery. Comparing to sero-negative control and convalescent COVID-19 patients with milder disease, the hospitalized patients had a dramatic reduction in the population of effector/activated Tregs (clusters T1_HLA-A_HLA-B and T5_HLA-DRA_S100A6), which are known important for protecting patients from tissue injury during respiratory virus infections including SARS-CoV-2(35). In contrast, the hospitalized patients had a relative expansion in TGF-β high cluster T4_KLF2_TGFB1 Tregs (**Figure 5A & B**). Interestingly, patients recovered from an asymptomatic infection showed a unique expansion of cluster T0_HLA-DR_CD74 relative to any of the severity groups. This cluster demonstrates the highest expression of class II HLA molecules (**Figure 5C, Supplementary Table S4**), which has previously been suggested to mark a specific highly suppressive Treg subset that primarily inhibits CD4^+^ helper T responses through contact-mediated suppression (36). In addition, our integrative analysis of single-cell Treg transcriptomes shows increased Cluster T0-HLA-DR_CD74 and Cluster T3-EEF1_IFITM1, but decreased Cluster T6-TCF7_GAS5, in asymptomatic and hospitalized patients. In contrast, Tregs in Clusters T2-ANXA1 and T7-KLRB1-LMS1 are decreased in all COVID groups (**Figure 5A)**.

**Figure 5.**
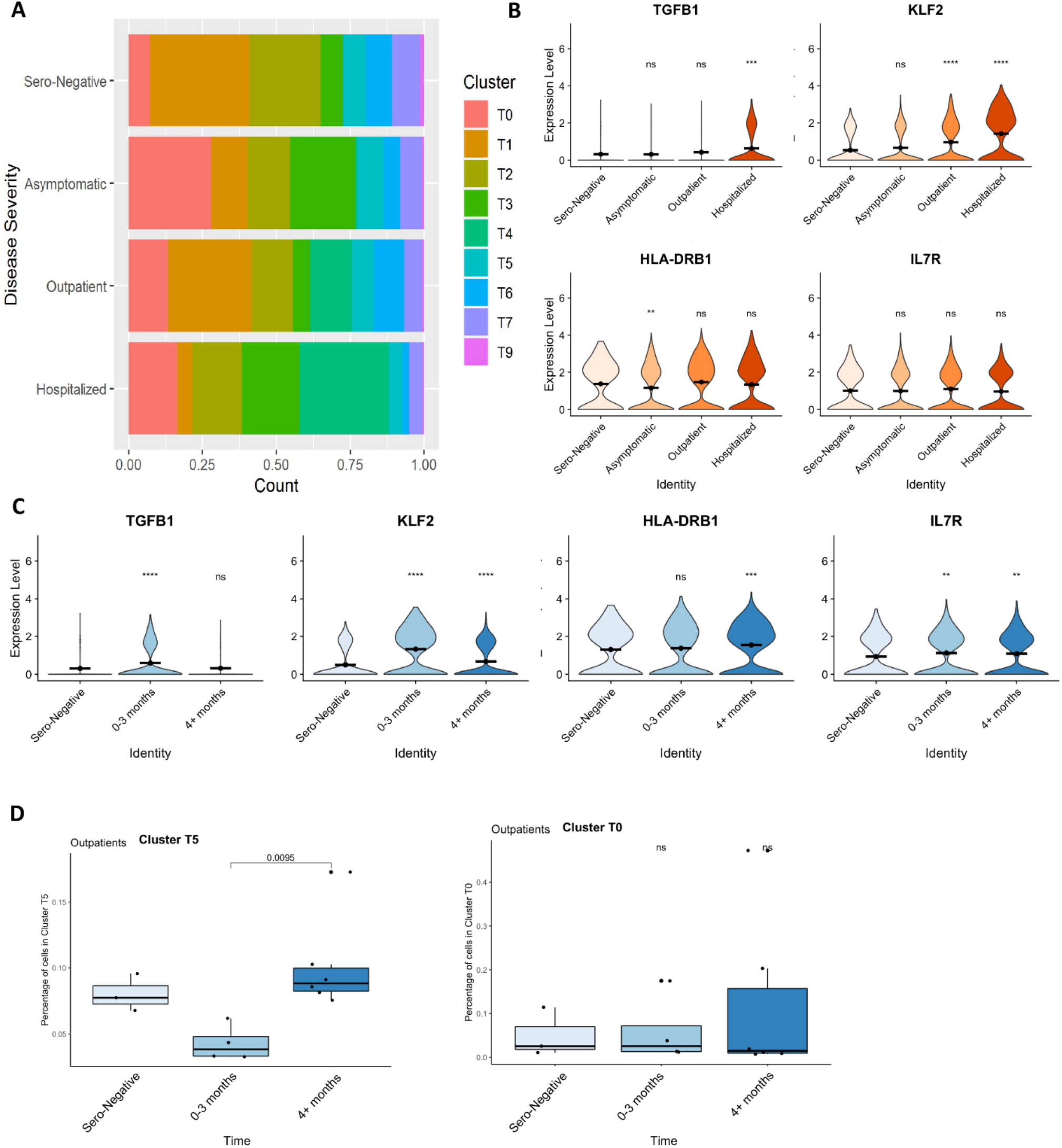
Association of Treg molecular signatures with COVID-19 disease severity and recovery. **(A)** Proportion of treg cells from each patient type in each cluster. Cluster T4 is only present among symptomatic COVID patients, not in asymptomatic or sero-negative patients. **(B)** Expression level of genes characteristic of cluster 4 separated by disease severity. KLF2 and TGFB1 appear particularly high in the most severe patients. **(C)** Expression of cluster T4 characteristic genes across time in outpatients and sero-negative controls. TGFB and KLF2 are most highly expressed 0-3 months post disease. **(D)** Cluster T5 (circulating Tregs) appears to decrease temporarily after COVID-19 illness in outpatients before recovering after 4+ months. Cluster T0 shows no changes after COVID-19 illness. B-C: Significance of gene expression difference in individual groups evaluated by Wilcoxon comparison of group-cell expression levels with Sero-Negative-cell expression levels. D: Boxplots indicate quartiles. Individual comparisons made using Wilcoxon signed rank test. Significance values: ns: p>0.05, * p<0.05, ** p<0.01, *** p<0.001, **** p<0.0001

Further analysis showed that both TGF-β1 and IL7R upregulation occurred during the early COVID-19 recovery phase (0-3 month), which gradually returned to normal during the later recovery phase whereas KLF2 increased and sustained over time in the Tregs from COVID-19 patients (**Figure 5C**).

We also noticed a significant decrease in Tregs of cluster T5_HLA-DRA_S100A6, which shows lower levels of FOXP3 in hospitalized patients, which is possibly due to the inflammation-induced downregulation of FOXP3 (**Figure 5A**). Similarly, the frequency of effector/activated Tregs with high expression of FOXP3 and STAT1 in cluster T5_HLA-DRA_S100A6 were dramatically reduced during the early recovery phase in COVID-19 outpatients, which returned to a comparable level similar to that in sero-negative controls 4 months after disease recovery. As expected, the intermediate Cluster T0_HLA-DR_CD74, which occupy one of the major Treg populations, did not show significant changes (**Figure 5D**). Collectively, our study identified a dynamic association of Treg frequency and their immune regulatory and tissue injury protective molecular signatures with disease severity identified in convalescent COVID-19 patients.

## Discussion

Our study enrolled 100 convalescent patients with different disease severities including asymptomatic, outpatient, hospitalized and ICU as well as healthy controls. All patients were enrolled at least three weeks after recovery from COVID-19 confirmed by negative in nucleic acid tests and complete disappearance in clinical symptoms. Their history with SARS-CoV-2 infection was all confirmed by the presence of abundant RBD-specific IgG in their circulating blood. We focused on the analysis of the monocytes, an innate immune cell population that are largely short-lived in the circulating blood, and the understudied regulatory T cells for dissecting the experienced immune responses of COVID-19 patients at the single-cell resolution.

Surprisingly, flow cytometry analysis revealed that the frequency of the short-lived monocyte and Treg populations display a sustained and dynamic association with disease severity in convalescent COVID-19 patients. Consistent with their dynamic frequency changes, we discovered several COVID-19-associated immune signatures such as the elevated IFN responsive genes in asymptomatic COVID-19 patients, which appears to be downregulated in patients with severe disease in particular the ICU patients. Many studies through single cell transcriptome analysis of PBMCs obtained during the active phase of the disease, have shown that the impairments in the IFN response is the key mediators of severe disease (13, 31). Our study here confirms that this dynamic association of IFN response with COVID-19 disease severity is sustained after full recovery. Interestingly, further longitudinal analysis showed that the elevated IFN responsive genes in monocytes could be sustained for more than 4 months of the studied time. In addition to IFN response, the altered HLA expression on myeloid cells during active SARS-CoV-2 infection, which is presumably due to the virus-induced inflammatory cytokines, have been identified(21, 37). Our study here demonstrated that this HLA high signature is also sustained after COVID-19 recovery. During the convalescent phase following COVID-19 infection, the immune system consists of two distinct pools of cells, COVID-19 experienced, such as memory T and B cells specific to SARS-CoV-2 antigens, and COVID-19 unexperienced cells. It has been well documented that the half-life of classical and non-classical monocytes in both mice and human is estimated at half-to 2.2 days(38-40). Therefore, in convalescent patients, the circulating monocytes and granulocytes are presumably unexperienced directly to SARS-CoV-2-specific antigens as well as the associated inflammatory environment during the acute phase of infection. Therefore, it is reasonable to speculate that the COVID-19-associated molecular signatures are largely inherited from their precursors and/or even the hematopoietic stem cells experienced with SARS-CoV-2 infection. Indeed, a recent study has suggested that the CD34^+^ hematopoietic stem/progenitor cells were primed toward megakaryopoiesis, accompanied by expanded megakaryocyte-committed progenitors and increased platelet activation by SARS-CoV-2 infection(20).

Tregs cells are known to modulate broad aspects of both innate and adaptive immune responses to maintain immune tolerance and global immunosuppression and consequently protect the host from autoimmunity(41). It has been well established that Tregs play a critical role in protecting tissue injury during microbial infections including influenza viruses and SARS-CoV-2(42, 43). Consistent with those observations note, our study here demonstrated that compared to sero-negative healthy controls, the Treg cell frequency was increased in COVID-19 patients with milder disease severity and this increase was not observed in hospitalized and ICU patients. As expected, our in-depth single cell transcriptome analysis of the sorted Treg populations identified 10 distinct clusters, confirming that human circulating Treg populations are highly heterogenous regardless with or without microbial infection(44-48). Interestingly, a Treg population (cluster T4_KLF2_TGFB1) that was characterized by elevated expression of TGF-β and KLF2, was only identified in some COVID-19 patients in particular the ICU patients, but not in sero-negative healthy controls. Further longitudinal analysis shows that the upregulation of both TGF-β and KLF2 sustained only during the first three months after full recovery, which was not observed in these convalescent COVID-19 patients 4 months after their full recovery. The clinical significance of this TGF-β and KLF2 high population is unclear. Since Treg-specific production of TGF-β is critical for a variety of immunologic roles, the fact that this population is only observed in patients recovering from COVID-19 suggests a possible role in COVID-19 pathogenesis and recovery. Consistent to our observation, it has been recently observed that SARS-CoV-2 in severe COVID-19 patients induces TGF-β expression in plasma blasts and possibly involved in IgA production for improving the lung mucosal immunity(49). A recent interesting study observed that Tregs from patients with severe disease produce a substantial amount of interleukin (IL)-6 and IL-18(50), which was not observed in COVID-19 patients in their recovery phase. It will be interesting to study how Treg frequencies and their COVID-19-associated molecular signatures dynamically change from active infection to recovery phase.

## Methods

### Human subject study and biosafety approvals for blood draws

All research activities with human blood specimens of pre-COVID-19, sero-negative (healthy) donors and convalescent COVID-19 patients were implemented under NIH guidelines for human subject studies and the protocols approved by the Northwestern University Institutional Review Board (STU00205299) as well as the Institutional Biosafety Committee for COVID-19 research. For collecting human blood specimens, patients and donors were recruited at Northwestern Memorial Hospital based on their availability and willingness to consent and participate in the research. Participants were compensated $20 and the blood was collected prior to COVID-19 vaccination.

All convalescent COVID-19 patients enrolled in our study meet the following criteria: testing positive for SARS-CoV-2, at least three weeks following recovery; some patients were also enrolled if they had a positive RBD-specific IgG in a community screening study. 20 mL of blood was drawn into EDTA tubes and transported on ice blocks to the Robert H. Lurie Comprehensive Cancer Center Flow Cytometry Core Facility at Northwestern University.

### Analysis of sera RBD-specific IgG levels in the plasma and its neutralization capacity

Plasma was centrifuged at 1,200 x g for 10 minutes at 4°C (brake off) after blood samples were diluted 1:1 with PBS during B cell isolation using RosetteSep Human B-Cell Enrichment Cocktail (Stem Cell Technologies). SARS-CoV-2 RBD-specific IgG ELISA tests with the plasma were performed in the lab of Dr. Alexis Demonbreun using enzyme-linked immunoassay (ELISA) as previously described (51). In brief, plasma was run in one or two duplicates and reported as the average. Results were normalized to the CR3022 antibody with known affinity to the receptor binding domain of SARS-CoV2 (Creative Biolabs, MRO-1214LC)(29, 52). Anti-RBD IgG concentration (μg/mL) was calculated from the 4PL regression of the CR3022 calibration curve. Neutralization assays with 10 μL or 80 μL of plasma were performed using HEK293 cells with stable expression of human ACE2. Commercially available SARS-CoV-2 spike receptor binding domain (RBD) (Raybiotech catalog no. 230-20407) was biotinylated using EZ-Link Sulfo-NHS-LC-Biotin (Thermo Scientific catalog no. 21335) and desalted with a Zeba quick spin column (Thermo Scientific catalog no. 89877). 5 μg biotinylated RBD was pre-coupled with 0.6 ul Streptavidin-AF647 (Invitrogen catalog no. S21374), used as RBD-647. 0.5 μL RBD-647 was then pre-incubated with PBS as a control or 80 μL patient-derived plasma for 45 minutes on ice, after which it was combined with 10^6^ HEK/ACE2 cells. These cells were incubated on ice for 45 minutes, washed and stained with DAPI. The levels of RBD-647 binding to HEK293/ACE2 cells were then assessed BD FACSAria Cell Sorter and analyzed using FlowJo v10software.

### Flow cytometry analysis of PBMCs and cell sorting of monocytes and Tregs

Total white blood cells from 1-2 mL freshly collected blood in EDTA tubes were enriched through ammonium chloride lysis (BD Bioscience, catalog no. 555899), followed by centrifugation (300x g for 10min). Supernatant was discarded, cells were washed twice using MACS buffer (Miltenyi, catalog no. 130-091-221); and pellet resuspended in 200uls MACS + Fc-block (ThermoFisher, catalog no. 14-9161-73). Cells were stained for surface antigens using fluorescence-conjugated monoclonal Abs specific to human CD45, CD3, CD4, CD127 and CD25 antibodies for Treg identification and sorting as well as CD45, CD3, CD64, CD14 and CD16 for monocyte characterization and sorting. Cells were also incubated with a patient-specific hashtag oligo (Biolegend TotalSeq-C0251-260 hashtag antibodies) as per manufacturer instruction. Post staining for 30 minutes on ice, cells were washed with MACS buffer and pellet resuspended in 1ml MACS buffer for sorting. Sorting was carried out on BD FACSAria SORP cell sorter housed in an independent negative pressure lab space in Baker BioProtect IV Biological Safety Cabinet. Sorted cell fractions were collected in MACS buffer and processed immediately for 10x library preparation. All of the sample preparation and cell sorting were carried out using BSL-2+ practices per Institutional Biosafety Committee approved protocols. Monocytes and Treg analyses with respective gating strategies are shown in Figure 1B. Detailed information of all fluorescent-conjugated monoclonal Abs used for this study are shown in supplemental Table S5.

### 10X library preparation

The concentration and viability of the single cell suspension were measured using a Nexcelom Cellometer Auto 2000 with ViaStain AOPI staining solution (Nexcelom, CS2-0106). Cells were loaded onto a 10x Chromium Controller for GEM generation followed by single cell library construction using 10x Chromium Next GEM Single Cell 5’ Library and gel bead kit v1.1 (10X Genomics, PN-1000165) according to manufacturer’s protocol. Quality control of the libraries was performed using an Invitrogen Qubit DNA high sensitivity kit (ThermoFisher Scientific, Q32851) and Agilent Bioanalyzer high sensitivity DNA kit (Agilent, 5067-4626). The libraries were pooled in equal molar ratio and sequenced on an Illumina HiSeq 4000 using sequencing parameters indicated by the manufacturer (Read 1: 26 cycles, i7 index: 8 cycles; Read 2: 91 cycles).

### scRNA-seq data acquisition and analysis

Data from scRNA-seq was demultiplexed and mapped to hg38 (refdata-gex-GRCh38-2020-A) using Cell Ranger software version 4.0.0 (10x genomics). Matrix files were analyzed using the Seurat R package (Seurat v4.0.3, R verson 4.1.0)(53). Libraries were loaded individually and log-normalized to correct for batch effects. Cells with mitochondrial reads >5% of total and fewer than 200 mRNAs were filtered out. Patient-specific hashtags were demultiplexed from each library. Libraries were then merged together and scaled across all samples. Monocytes and Treg clusters were identified by expression of CD14, CD16, FOXP3 and CD25. Monocyte and Treg clusters were split from one another and analyzed separately. Principal component analysis was performed on each data set, and Uniform Manifold Approximation and Projections (UMAP) were made from the top 10 principal components. Positive markers for each cluster were identified with a minimum log-FC threshold of 0.25. Statistically significant markers with the highest log-FC were used in the creation of heatmaps.

### Statistical analyses

Kruskal-Wallis tests were used to identify overall significant differences at the population level. Wilcoxon signed-rank tests were used to identify significant differences between a sub-group and the overall population (Figures 2C and 4C) or to compare individual subgroups against one another (Figures 1, 3 and 5). All statistics were computed on R v 4.1.0

### Data availability

The single cell RNA sequences are to be deposited pending acceptance at dbGaP.

## Acknowledgments

We are thankful to the team of Northwestern COVID-19 Antibody and Cancer Collaborative Group and advisory members, especially Drs. Alfred L. George Jr., Richard D’Aquila, Leonidas C. Platanias, Rex L. Chishom, and William A. Muller for their scientific input and resourceful support for the project. The work was partially funded by Chicago Biomedical Consortium Accelerator Award A-017 (H.L. and D.F.), Northwestern University Feinberg School of Medicine Emerging and Re-emerging Pathogens Program (EREPP) (H.L.), Department of Pharmacology Start-up fund (H.L.), and the R.H. Lurie Comprehensive Cancer Center Blood Biobank fund and Northwestern University NUCATs (M.I.). We gratefully acknowledge the support from the NUseq Core Facility. This work was supported by the Northwestern University RHLCCC Flow Cytometry Facility and a Cancer Center Support Grant (NCI CA060553). Flow Cytometry Cell Sorting was performed on a BD FACSAria SORP system and BD FACSymphony S6 SORP system, purchased through the support of NIH 1S10OD011996-01 and 1S10OD026814-01.

## Author contributions statement

A.D.H, and S.E.W. designed and led the bench experiments, analyzed data, prepared figures, and contributed to writing. S.S., S.C., H.F.M, M.J.S., C.M., X.W., L.E., N.K.D., J.W., P.J.M., L.J.S., C.M.W., C.O., Y.J., P.D., and N.R.W. provided technical support or conducted bench experiments, and analyzed data. Y.L., A.R.D., and M.G.I. provided critical resources and supervised the research project. D.F., and H.L designed experiments, analyzed data, wrote the manuscript, and supervised the work.

## Competing interest statement

Northwestern University and H. L., D. F., L. E., A. D. H., and N. K. D. hold issued and/or provisional patents in the area of exosome therapeutics and COVID-19 therapeutics. H.L., D. F., and A.D.H are scientific co-founders in ExoMira Medicine Inc.

**Supplementary Table S1.**

| **Monocytes** |  |  |
| --- | --- | --- |
|  | **Patients (number)** | **Cells (number)** |
| Total | 50 | 18165 |
| **By Disease Severity** |  |  |
| Sero-Negative | 3 | 593 |
| Asymptomatic | 11 | 2379 |
| Outpatient | 25 | 11877 |
| Hospitalized | 7 | 1707 |
| ICU | 4 | 1609 |
| **By Time** |  |  |
| Sero-Negative | 3 | 593 |
| 0-1 months | 6 | 2624 |
| 2-3 months | 16 | 6448 |
| 4+ months | 19 | 5616 |
| Unknown | 6 | 2884 |

**Supplementary Table S2.**

| **Fig 3B and 3D**<br>**Outpatients/SN Only** | **Patients (number)** | **Cells (number)** |
| --- | --- | --- |
| Sero-Negative | 3 | 593 |
| 0-1 Months | 6 | 2624 |
| 2-3 Months | 7 | 4463 |
| 4+ Months | 10 | 2899 |

**Supplementary Table S3.**

| **Tregs** |  |  |
| --- | --- | --- |
|  | **Patients (number)** | **Cells (number)** |
| Total | 27 | 7178 |
| **By Disease Severity** |  |  |
| Sero-Negative | 3 | 483 |
| Asymptomatic | 10 | 2289 |
| Outpatient | 10 | 2749 |
| Hospitalized | 4 | 1057 |
| **By Time** |  |  |
| Sero-Negative | 3 | 483 |
| 0-3 months | 11 | 2960 |
| 4+ months | 9 | 2620 |
| Unknown | 4 | 1115 |

**Supplementary Table S4.**

| **Fig 5c**<br>**Outpatients/SN Only** | **Patients**<br>**(number)** | **Cells (number)** |
| --- | --- | --- |
| Sero-Negative | 3 | 483 |
| 0-3 Months | 4 | 1177 |
| 4+ Months | 6 | 1572 |

**Supplementary Table S5.**

| **Antibody** | **Fluorophore** | **Clone** | **Cat # (Company)** |
| --- | --- | --- | --- |
| **CD3** | APC-e780 | SK7 | 47-0036-41 (eBioscience) |
| **CD4** | BUV737 | SK3 | 612749 (BD Biosciences) |
| **CD25** | BUV395 | 2A3 | 564034 (BD Biosciences) |
| **CD127** | BV650 | A01905 | 351325 (Biolegend) |
| **CD14** | PE | MOP9 | 562691 (BD Biosciences) |
| **CD16** | APC | 3G8 | 561248 (BD Biosciences) |
| **CD64** | PE-Cy7 | 10.1 | 561191 (BD Biosciences) |
| **Ace2** | Alexa 488 | 535919 | FAB9332G (R&D) |
| **CD45** | BV510 | HI30 | 563204 (BD Biosciences) |
| **DAPI** | DAPI | - | D1306 (ThermoFisher) |

## Figures and legends

**Figure S1:**
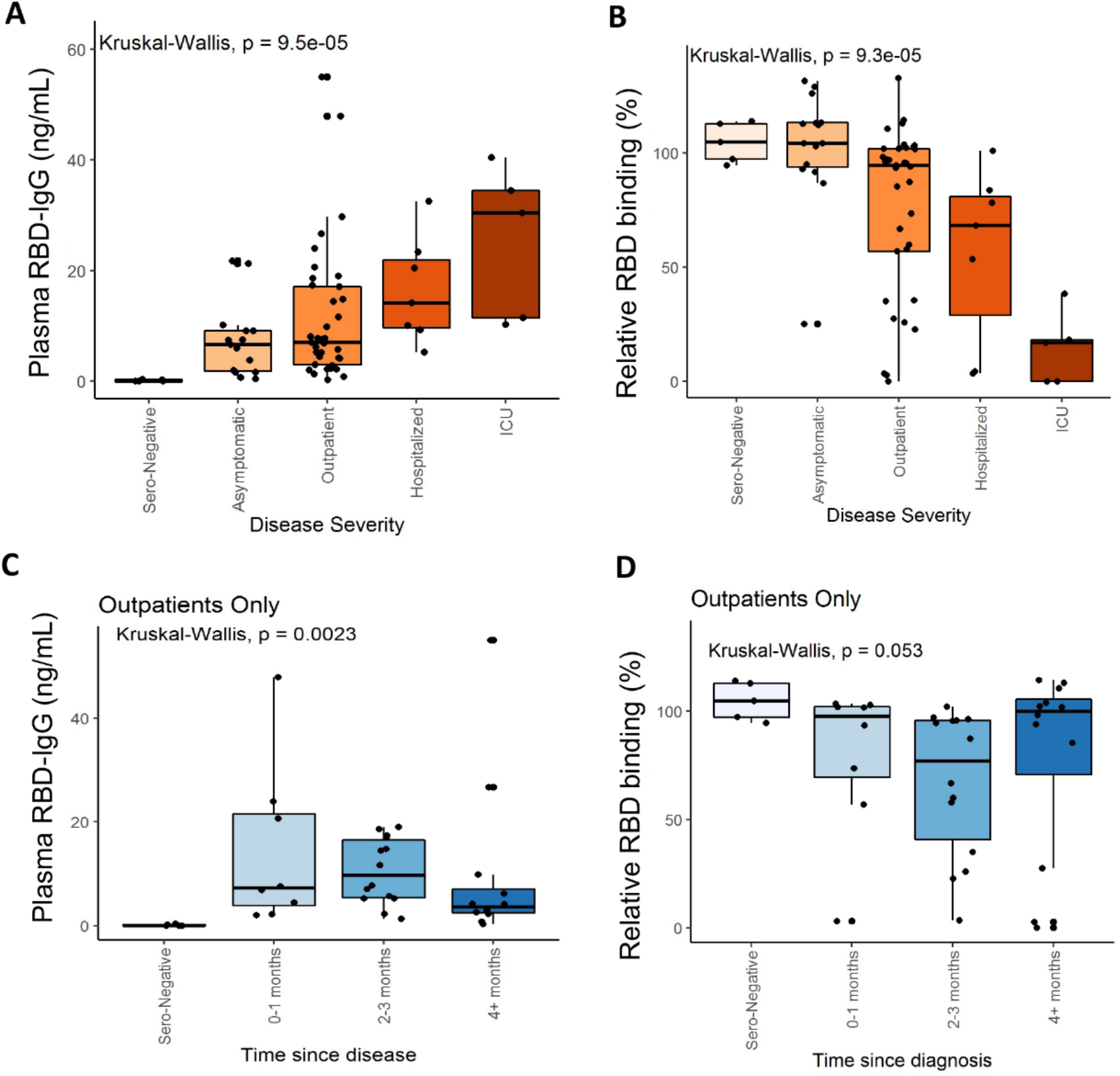
IgG levels and blocking capacity in convalescent patient plasma. **(A)** Schematic showing the treatment of patient-derived blood samples **(B)** Anti-SARS-CoV-2-spike protein IgG levels in patients, separated by disease severity. **(C)** Ability of patient-derived plasma to inhibit binding of AlexaFluor-647 tagged SARS-CoV-2-spike receptor binding domain (RBD) to ACE2-expressing HEK-293 cells. Normalized to a no-plasma (PBS only) control. Higher values indicate less inhibition of the binding. **(D)** IgG levels as in B, but outpatient samples only separated by time since disease. **(E)** RBD binding capacity as in (C) but outpatients only and separated by time since disease. Boxplots in B-E indicate quartiles. Significance evaluated through Kruskal-Wallis test: ns: p>0.05, * p<0.05, ** p<0.01, *** p<0.001, **** p<0.0001.

## Notes

### Competing Interest Statement

The authors have declared no competing interest.

